# Millimeter-resolution mapping of citrate exuded from soil grown roots using a novel, low-invasive sampling technique

**DOI:** 10.1101/2020.12.09.417550

**Authors:** Raphael Tiziani, Markus Puschenreiter, Erik Smolders, Tanja Mimmo, José Carlos Herrera, Stefano Cesco, Jakob Santner

**Author notes:** Corresponding authors: **Raphael Tiziani**, Faculty of Science and Technology, Free University of Bozen-Bolzano, Italy Piazza Università 5, I-39100 Bolzano, **Jakob Santner**, Institute of Agronomy, Department of Crop Sciences, University of Natural Resources and Life Science, Vienna, Austria, Konrad-Lorenz-Straße 24, AUT-3430 Tulln an der Donau.

## Abstract

The reliable sampling of root exudates in soil grown plants is experimentally challenging. This study aimed at developing a citrate sampling and mapping technique with millimetre-resolution using DGT (diffusive gradients in thin films) ZrOH binding gels. Citrate adsorption kinetics, DGT capacity and stability of ZrOH gels were evaluated. ZrOH gels were applied to generate 2D maps of citrate exuded by white lupin roots grown in rhizotrosn in a phosphorus deficient soil. Citrate was adsorbed quantitatively and rapidly by the ZrOH gels, these gels can be stored after sampling for several weeks prior to analysis. The DGT capacity of the ZrOH gel for citrate depends on the ionic strength and the pH of the soil solution but was suitable for citrate sampling. 2D citrate maps of rhizotron grown plants have been generated for the first time at a millimetre resolution to measure an illustrated plant response to P fertilization. DGT-based citrate sampling is suitable for studying the root exudation in soil environments, at unprecedented spatial resolution. By changing binding material, the technique is also applicable to other exudate classes and might be used for the evaluation of whole root exudation crucial in specific cultivar breeding.

**Highlight:** We present a novel, reliable, easy to use, non-destructive citrate sampling- and two-dimensional high-resolution imaging technique for soil grown plant roots.

## Introduction

Root exudates comprise a myriad of soluble organic compounds excreted by plant roots in the rhizosphere in order to improve its edaphic conditions. The share of organic carbon (C) exuded differs greatly depending on the plant species and/or even on the genotype, the age, the chemical, physical and biological soil characteristics and biotic or abiotic stresses (Bais *et al*., 2006; Jones *et al*., 2009; Badri and Vivanco, 2009; Meier *et al*., 2013; Canarini *et al*., 2016; Tückmantel *et al*., 2017; Nakayama and Tateno, 2018; Miller *et al*., 2019). The majority of exudates are primary metabolites including organic acids, carbohydrates and amino acids, followed by secondary metabolites such as phenols, vitamins, glucosinolates, plant hormones, but also enzymes and polysaccharides (Jones *et al*., 2009; Badri and Vivanco, 2009). Root exudates shape the biological, chemical, biochemical, and physical properties of the rhizosphere (Traoré *et al*., 2000; Shi *et al*., 2011*b*; Steinauer *et al*., 2016; Naveed *et al*., 2017; Baumert *et al*., 2018).

Organic acids belong to the most important root exudates due to their impact on nutrient dynamics. Particularly, citrate plays a huge role in root phosphorus (P) acquisition (López-Bucio *et al*., 2000; Lyu *et al*., 2016; Giles *et al*., 2018; Wang and Lambers, 2019) by solubilizing P bound to iron(Fe) or aluminium(Al) (hydr)oxides (Otani *et al*., 1996). Abiotic stresses, such as P deficiency, trigger citrate exudation (Meyer *et al*., 2010; Mora-Macías *et al*., 2017). The knowledge about the quality, quantity and spatial allocation of citrate is of immense importance to understand rhizosphere dynamics and mechanisms, but it may also help to optimize agricultural productivity and quality. However, the reliable quantification and localization of exudates like citrate released from roots growing in soil is a very challenging task.

Traditional root exudate sampling methods can be divided into two main categories: i) sampling from soil-grown plants or ii) from hydroponically grown plants. Collecting exudates directly from rhizosphere soil is very difficult for several reasons: restricted access to the root, very low reproducibility, rapid microbial degradation of exudates and sorption to the solid soil phase (Oburger and Jones, 2018; Vives-Peris *et al*., 2019). Growing plants first in soil and then collecting the exudates in an aqueous solution may be a good alternative. However, the needed soil removal can damage the roots and alter the exudation profile (Oburger and Jones, 2018). Moreover, other soil-based methods include the use of root exudate collection tools (SOIL-REC) (Oburger *et al*., 2013), RHIZOtests (Chaignon *et al*., 2002) or micro-suction cups (Puschenreiter *et al*., 2005). Sampling exudates from individual roots in soil can be achieved by using specialized resins (Shi *et al*., 2011*a*), filter paper (Neumann *et al*., 2014), agar sheets (Hoffland *et al*., 1989) or excavation/cuvette-hydroponic-sampling (Phillips *et al*., 2008). Most of the mentioned techniques are very complex like the SOIL-REC or disturb roots greatly like the excavation sampling. Therefore, most studies about root exudates are still performed in hydroponic-based both plant-growth and sampling systems (Vives-Peris *et al*., 2019). The advantages are a higher reproducibility, possible sterility, easy access to soil-free roots and the ability to easily control environmental conditions (Vranová *et al*., 2013; Tiziani *et al*., 2020). To sample the exudates hydroponically, the nutrient solution is replaced by a collecting solution for a certain amount of time (Valentinuzzi *et al*., 2015*a*). Exudates from single root segments can be sampled by trapping them into small chambers, placing the roots onto adsorption resin agar or by the application of agar sheets (Marschner *et al*., 1987; Hoffland *et al*., 1989). Plant development, physiology and root exudation are however affected by hydroponic growing systems since they are highly artificial environments. Moreover, hydroponic sampling is usually limited to greenhouses/climate chambers and their application causes often mechanical stress, which further affects exudation patterns (Oburger and Jones, 2018; Vives-Peris *et al*., 2019). Thus, results obtained in hydroponics cannot directly be transferred to soil-plant systems raising questions about the ecological relevance of the obtained information (Oburger and Jones, 2018).

So far, no unbiased exudate sampling method exists and each of the methods has its shortcomings, it is logical that soil-based systems are preferred over solution for providing environmentally relevant data (Oburger and Jones, 2018; Wang and Lambers, 2019). Therefore, there is still the urgent necessity to develop reliable and meaningful soil-based root exudation sampling methods to improve the knowledge about rhizosphere processes (Wang and Lambers, 2019). Diffusive gradients in thin films (DGT) may offer a yet unexplored possibility because it is non-invasive and allows to concentrate and preserved the exudates. DGT is a passive sampling technique originally developed for applications in aquatic chemistry (Davison and Zhang, 1994). Today, DGT is a well-established method used for the measurement of labile inorganic and organic substances in natural waters, soils and sediments and for two-dimensional high-resolution chemical imaging (Santner *et al*., 2015; Davison, 2016). During the DGT application, a gel is deployed on the surface of the environment (here: a rooted soil section), that gel contains a binding agent typical for the analyte, the labile analyte diffuses from the environmental matrix through a thin diffusion layer, usually composed of a protective membrane and a thin polyacrylamide gel, towards a binding gel containing to the analyte-binding phase, e.g. zirconium (hydro)oxide (ZrOH), ferrihydrite or Chelex 100 (Santner *et al*., 2015). The gel is subsequently removed and analysed in an either destructive way by extracting sections of the gel or by mapping the analytes in 2D with surface detection methods. This DGT technique has been successfully applied for the sampling of inorganic substances from the rhizosphere (Santner *et al*., 2012; Valentinuzzi *et al*., 2015*b*). The application of DGT for exudate sampling with an appropriate binding material has the potential (1) to sample and concentrate exudates directly from individual soil-grown plant roots without damaging the root (2) to protect the exudate from mineralization during sampling by immobilising it on the binding material and (3) to allow for localisation of citrate exudation along and perpendicular to the root axis at millimetre resolution.

This study was set up to test the suitability of a DGT-based exudate sampling and mapping technique using ZrOH binding gels (Guan *et al*., 2015) and established DGT rhizosphere sampling protocols (Wagner *et al*., 2020). Zirconia based adsorbents have a great capacity and affinity to bind carboxylates (Stern *et al*., 2014; Jia *et al*., 2019) and was the selected binding agent here. The exudate binding properties were tested. Effects of gel storage on citrate stability were tested in the absence or presence of soil to identify potential citrate decomposition after sampling. Finally, a case study was performed by mapping citrate concentrations in the rhizosphere of white lupin (*Lupinus albus* L.) in response to P fertilisation in a P-deficient soil.

## Materials and Methods

### General laboratory procedures and materials

All high purity chemical standards were purchased from Sigma-Aldrich Germany. High performance liquid chromatography (HPLC) grade chemicals were used for the ion chromatography (IC) analysis. Laboratory water type I (0.055 μS cm^−1^; TKAGenPure, Niederelbert, DE) was utilized for preparing all experimental solutions and for all analyses. All citrate solutions, except the ones used for testing citrate degradation on the ZrOH DGT gel, were prepared with 3 mmol L^−1^ sodium azide (NaN_3_) to prevent microbial degradation of citrate. All experiments were conducted with three replicates (n = 3).

The soil used in this study originated from the mineral surface layer (0-20 cm) of a ferralsol from Behenjy, Madagascar, which was chosen because of its particularly low P bioavailability. This soil had a pH of 4.17 (CaCl_2_), a soil organic C content of 3.3 g kg^−1^, a total P content of 629 mg kg^−1^ (*aqua regia*) and 29.3 mg kg^−1^ ammonium-oxalate-extractable P. The water holding capacity (WHC) was 60 % (w/w). The extent of P deficiency in this soil is so strong that plant growth fails in absence of P fertiliser (De Bauw *et al*., 2020).

Prior to the DGT and plant experiments, the soil was limed with 0.92 g CaO kg^−1^ and fertilized with the following salts (mg kg^−1^): NH_4_NO_3_ 285, KH_2_PO_4_ 0 and 1098.5 (for 0P and 250P respectively, the latter being the 250 mg P kg^−1^ soil equivalent), KCl 63.5, MgSO_4_ × 7 H_2_O 210, FeCl_3_ × 6 H_2_O 4, MnCl_2_ × 2 H_2_O 2.45, ZnCl_2_ 2.26, CuCl_2_ × 2 H_2_O 2.23, H_3_BO_3_ 0.47 and Na_2_MoO_4_ × 2 H_2_O 0.37. Liming resulted in a pH of 5.25 (CaCl_2_). The ionic strength of a saturated-paste extract sampled 45 days after liming and fertilization, at the time of citrate sampling from roots using DGT gels, was 19.6 mmol L^−1^. Additional soil chemical characteristics, as well as the liming and fertilization procedure, are given in Supplementary Table 1 and Supplementary Information (SI), respectively.

**Table 1.**
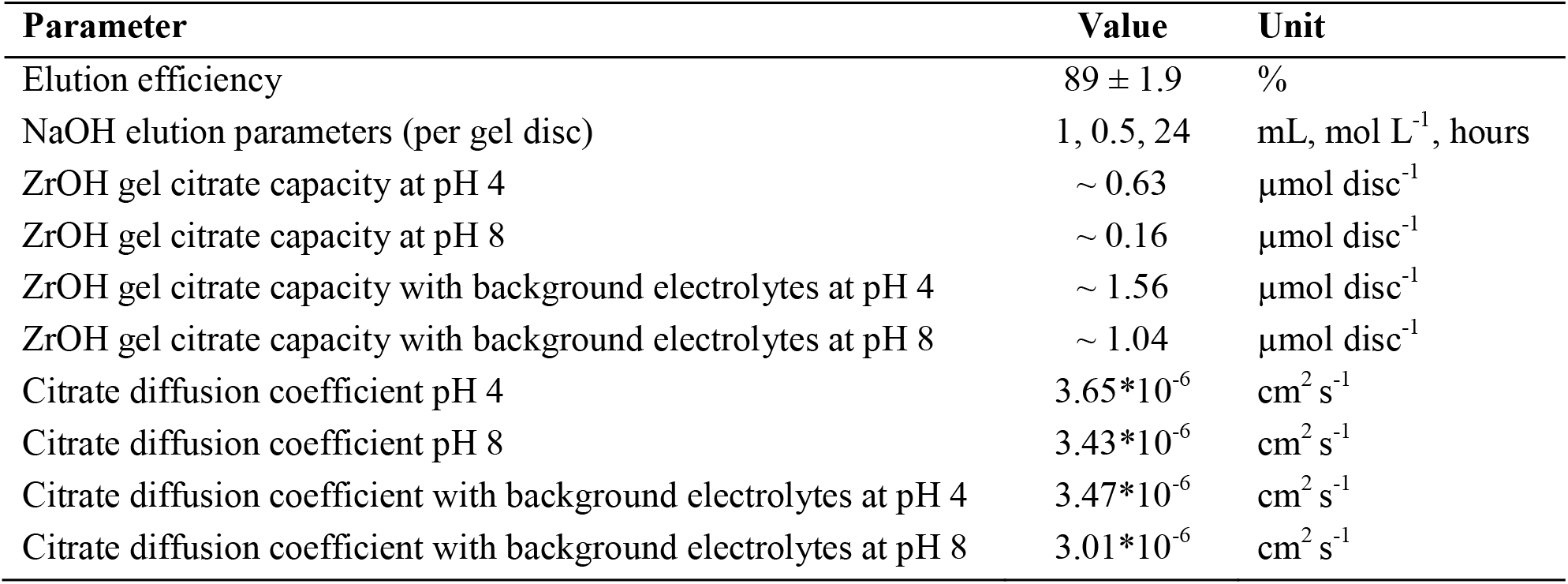
Chemical characteristics of ZrOH gels for citrate sampling.

| Parameter | Value | Unit |
| --- | --- | --- |
| Elution efficiency | $89 \pm 1.9$ | % |
| NaOH elution parameters (per gel disc) | 1, 0.5, 24 | mL, mol L<sup>-1</sup>, hours |
| ZrOH gel citrate capacity at pH 4 | ~ 0.63 | μmol disc<sup>-1</sup> |
| ZrOH gel citrate capacity at pH 8 | ~ 0.16 | μmol disc<sup>-1</sup> |
| ZrOH gel citrate capacity with background electrolytes at pH 4 | ~ 1.56 | μmol disc<sup>-1</sup> |
| ZrOH gel citrate capacity with background electrolytes at pH 8 | ~ 1.04 | μmol disc<sup>-1</sup> |
| Citrate diffusion coefficient pH 4 | $3.65 \cdot 10^{-6}$ | cm<sup>2</sup> s<sup>-1</sup> |
| Citrate diffusion coefficient pH 8 | $3.43 \cdot 10^{-6}$ | cm<sup>2</sup> s<sup>-1</sup> |
| Citrate diffusion coefficient with background electrolytes at pH 4 | $3.47 \cdot 10^{-6}$ | cm<sup>2</sup> s<sup>-1</sup> |
| Citrate diffusion coefficient with background electrolytes at pH 8 | $3.01 \cdot 10^{-6}$ | cm<sup>2</sup> s<sup>-1</sup> |

### Diffusive and binding gel preparation

Due to having the highest reported capacity for binding phosphate (65 µg cm^−2^, equivalent to 0.68 µmol cm^−2^), and because the established elution procedure for phosphate in 0.5 mol L^−1^ NaOH, expected to be suitable and convenient for eluting citrate as well, the precipitated ZrOH DGT binding gel of Guan *et al*. (2015) was chosen as candidate binding gel for citrate. As will be shown in the Results and Discussion sections, this gel proved to be well suited for DGT-based citrate sampling.

Polyacrylamide diffusive gel sheets (0.4 and 0.8 mm), as well as ZrOH gel sheets (0.4 mm), were prepared. Briefly, the gel solution made of 15% acrylamide, 0.3% DGT gel cross-linker (DGT Research Ltd, Skerlmorlie, Bay Horse Rd, Quernmore, Lancaster LA2 QQJ, UK), 0.7 % ammonium persulphate (10%, APS) and 0.25% of N,N,N’N’-tetramethylethylenediamine (TEMED, 99%) was cast between two glass plates (either for 0.4 or 0.8 mm gels) and immediately put in an oven at 45°C for 45 min. After the gels were completely set, they were soaked successively three times in at least 2 L of water for washing off remaining reagents and stored in a 0.01 mol L^−1^ NaNO_3_ solution. The 0.4 mm gels were used for the making of ZrOH gels. Briefly, 3.22g ZrOCl_2_ x 8 H_2_O was dissolved in ∼40 mL water. Four 0.4 mm diffusive gel sheets have been added into the ZrOCl_2_ x 8 H_2_O solution which was filled up to 100 mL to let the gels soak for at least two hours (0.18 mol L^−1^ ZrOCl_2_ x 8 H_2_O final concentration). Afterwards, each gel was transferred to 100 mL 0.05 mol L^−1^ 2-(N-morpholino)-ethanesulfonic acid (MES) at pH 6.70 while stirring gently for the first 60 seconds to obtain a homogenous precipitation. The gels were shaken gently for 40 min to allow complete precipitation of the zirconium (Zr). The ZrOH gels were washed with water and stored in 0.03 mol L^−1^ NaNO_3_ (Zhang and Davison, 1995; Zhang *et al*., 1998; Luo *et al*., 2010; Guan *et al*., 2015). Diffusive gels and ZrOH gels were cut in discs of 2.5 cm diameter for the necessary operations. For the application to sample citrate, ZrOH gels were cut into *square* sheets of different size (*e*.*g*. 3*2 cm).

### Characterization of ZrOH gel for citrate sampling

The ZrOH binding gel was tested for its citrate uptake and elution characteristics, for its citrate capacity, and for the effects of pH and ionic strength on citrate uptake. Citrate binding to ZrOH might be affected by the ionic composition of the sampled porewater, either by ion competition effects, or by changes in the net charge on the ZrOH phase due to sorption of non-target anions and cations. Most of the experiments described in the following were done in aqueous solutions with 3 mmol L^−1^ NaN_3_ as a biocidal treatment to avoid citrate decomposition. Selected experiments were done with a high, but realistic (Blume *et al*., 2010) concentration of background electrolytes with following concentrations (Sigma Aldrich, DE, mmol L^−1^): NO_3_^−^ 2.96, PO_4_ ^3−^ 0.007, K^+^ 0.68, Ca^2+^ 1.48, Mg^2+^ 1.01, SO_4_ ^2−^ 1.16, Al^3+^ 0.09, Cl^−^ 2.58, HCO_3_ ^−^ 1.31 and NaN_3_ 3.00, corresponding to an ionic strength of 15.8 mmol L^−1^ for investigating citrate binding to the ZrOH gel in low and high ion-background situations.

As a standard, gel characterization was done at the ambient pH of the experimental solutions (close to pH 5.6 of pure water). However, some tests were done with solution pH values of pH 4, buffered using 2 mmol L^−1^ benzoic acid or pH 8, buffered using 2 mmol L^−1^ 3-(N-morpholino)propanesulfonic acid (MOPS), to encompass the pH range that may be encountered in many soils. The specific experimental conditions are mentioned for each experiment.

#### Citrate elution efficiency and uptake kinetics

In order to obtain the most efficient citrate elution procedure, elution efficiency tests with different volumes and concentrations of NaOH were carried out. For the volume test, ZrOH gel discs (2.5 cm diameter) were loaded with citrate by immersion in a 10 mL solution containing 27.1 µg (0.11 µmol) citrate and 3 mmol L^−1^ NaN_3_ for 4 hours while shaking gently. The elution was subsequently carried out in 1, 2, 5 and 10 mL 0.5 mol L^−1^ NaOH for 24 hours while shaking gently. In a similar experiment ZrOH gel discs loaded with 25.4 µg (0.13 µmol) citrate were eluted in 1 mL of solutions containing 0.3, 0.4, 0.5, 0.6 or 0.7 mol L^−1^ NaOH for 24 h. The original immersion solutions, the immersion solutions after exposure to the gel and the eluates were analysed for citrate. The experiment was conducted with three replicates.

The most efficient elution (89%) was achieved when eluting in 1 mL 0.5 mol L^−1^ NaOH for a ZrOH disc with a volume of 0.196 cm^3^ for 24 hours (see Results). Each subsequent citrate elution was carried out using these parameters, maintaining the given ratio of eluent volume to gel disc volume constant. This was relevant and necessary also for the small gel pieces obtained in the plant experiment for mapping citrate exuded from white lupin cluster roots.

Citrate uptake kinetics on ZrOH gels were determined by immersion of 0.4 mm thick ZrOH gel discs in 10 mL of solutions containing 27.7 µg (0.14 µmol) citrate for 1, 5, 10, 30, 60, 120, 300 and 1440 minutes while shaking gently to ensure that the gel was in full contact with the solution. After the uptake period, the ZrOH gel was removed from the immersion solution and transferred to 1 mL of 0.5 mol L^−1^ NaOH for 24 h for eluting the citrate from the ZrOH binding phase. The original immersion solutions, the immersion solutions after exposure to the gel and the eluates were analysed for citrate. The experiment was conducted with three replicates.

#### Capacity of precipitated ZrOH gel for citrate binding

The capacity of ZrOH gels for binding citrate was determined using standard solution DGT samplers (DGT Research Ltd., Lancaster, UK), which allow for determining the gel binding capacity up to which the binding gel acts as a zero sink for a given solute. DGT samplers consist of a 0.4 mm thick DGT binding gel disc of 2.5 cm diameter (precipitated ZrOH in this study), overlain by a 0.8 mm thick diffusive gel disc and a protective membrane disc (polyethersulfone, 0.45 μm pore size, 0.15 mm thick, Sartorius Stedim, Göttingen, DE), encased by a plastic sampler housing (see Zhang & Davison (1995) for a detailed description). When immersed into a simple synthetic solution, DGT provides a measurement of the time-averaged concentration, *c*_DGT_, at the sampler-solution interface:

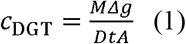

For determining *c*_DGT_, a few parameters on the sampler geometry and the analyte diffusion have to be known: *M* is the mass of citrate taken up by the sampler which is measured after eluting the binding gel, Δ*g* is the thickness of the diffusive layer (diffusive gel + protective membrane), *D* is the citrate diffusion coefficient in the diffusive gel, *t* is the deployment time and *A* is the surface area of the DGT sampler exposed to the sampling solution (A=3.14 cm^2^). If the DGT sampler acts as a zero sink for the analyte, *i*.*e*. binds the total amount of analyte that diffuses towards the binding gel, *c*_DGT_ equals *c*_soln_, the solution concentration. The capacity of a DGT gel to bind an analyte species is conventionally defined as the solute amount that can be bound while satisfying *c*_DGT_ = *c*_soln_. Above a certain amount of solute bound, *c*_DGT_ < *c*_soln_, as the gel ceases to bind the analyte immediately and quantitatively, and therefore ceases to act as infinity sink (Zhang and Davison, 1995). While the DGT binding capacity is not a measure of the total amount of analyte that can be bound by the binding gel, it serves as a conservative estimate up to which binding gel loading DGT exerts its infinite sink function. For this reason, the DGT binding capacity was determined in this work.

Three DGT samplers were deployed in 2 L of well-stirred solutions containing citrate at concentrations between 1.26 and 65.2 mg L^−1^ (6.55 and 339 µmol L^−1^) and 3 mmol L^−1^ NaN_3_ as a biocidal treatment for 30 hours. The experiment was performed at pH 4 and 8, as well as with and without background electrolytes, as described previously. The experiments were conducted with three replicates, *i*.*e*. three samplers in one 2L box. The procedure for determining the diffusion coefficient of citrate in the diffusive gel is given in the SI.

#### Citrate stability on ZrOH gels in the presence or absence of soil as potential citrate source

First, effects of cold storage on citrate loaded gels were tested. ZrOH gel discs were loaded with 8.6 µg (0.04 µmol) citrate by immersion in 10 mL of a solution without NaN_3_ for 4 hours. The discs were put into an airtight plastic bag with some drops of water for avoiding desiccation and stored in the fridge at 4°C. Gels were removed after 1, 3, 7, 10, 14, 29 and 41 days for citrate elution and analysis. The experiments were performed with three replicates.

Secondly, the effect of soil as a citrate source or a sink (degradation) were tested. The previously described 0P soil was either sterilized or not by adding 100 mg HgCl_2_ kg^−1^ (0.37 mmol kg^−1^) (Trevors, 1996). The appropriate amount of HgCl_2_ was diluted in 10 mL water, added to the correct soil mass and mixed well. The untreated soil received an addition of 10 mL water containing no HgCl_2_. Both soil subsamples were then moistened to 70% WHC, mixed well for 10 minutes and equilibrated for 3 days. Water content was increased to 80% WHC two hours prior to the DGT application. DGT probes loaded with 70.0 µg (0.37 µmol) citrate were exposed to unsterilized soil, while non-citrate containing DGT samplers where exposed to unsterilized and sterilized soil samples. The soil was applied onto DGT samplers, making sure that no air bubbles were entrapped between the DGT probes and the soil, and incubated for 6, 12, 24 and 48 hours at 24°C. Therefore, the samplers were put into an airtight plastic container together with wet paper to prevent soil drying. After each sampling period, the soil was removed by rinsing the probe with water, followed by the removal of the ZrOH gel from the sampler with subsequent elution and citrate analysis. The experiments were performed with three replicates.

### Case study: Mapping citrate exudation of *Lupinus albus* L. roots

#### Plant growth parameters

*Lupinus albus* L. (white lupin) plants were grown in rhizotrons with the same design as described in Santner *et al*. (2012). Rhizotrons are flat growth containers with detachable front plates. They were chosen to permit physical access to the roots and rhizosphere soil, as the roots grow in the surface layer of the contained soil along the removable front plate. The rhizotrons therefore allow the application of the ZrOH gels for citrate sampling.

The rhizotrons were made from perspex and have internal dimensions of 10 cm × 1.5 cm × 40 cm (w*h*l). Fourteen holes in the front plate allowed uniform irrigation during plant growth and permited saturating the soil with water before the ZrOH application. They were packed with the 0 and 250P soil (described above) to a uniform bulk density of 1.15 kg L^−1^. A layer of 10 µm thick polycarbonate membrane (Nucleopore, Whatman, Maidstone, UK) was placed between the soil and the removable front plate. In addition, a protective plastic foil was put between the polycarbonate membrane and the front plate. Rhizotrons were moistened to 50% WHC and kept constant during plant growth.

White lupin seeds were germinated in moist tissue paper for seven days. Once germinated, one seedling was transferred to each rhizotron. Aluminium foil was wrapped around the rhizotrons to prevent photo-chemical reduction phenomena in the rhizosphere and biofilm formation. For plant growth the soil was kept at 50% WHC. Rhizotrons were kept in a climate chamber with a day/night cycle of 14/10h and a day/night temperature of 20/15 °C.

#### Citrate sampling from white lupin roots

The sampling protocol has been published on the protocols.io website (doi: dx.doi.org/10.17504/protocols.io.bqgnmtve). DGT citrate sampling from white lupin roots was performed 45 days after transplanting when the plants had developed mature cluster roots (Fig. 1). For the sampling, soil water content was progressively brought to saturation over two hours. For the ZrOH gel application, the rhizotrons were laid flat onto a stand with the removable front plate facing upwards and were opened.

**Figure 1.**
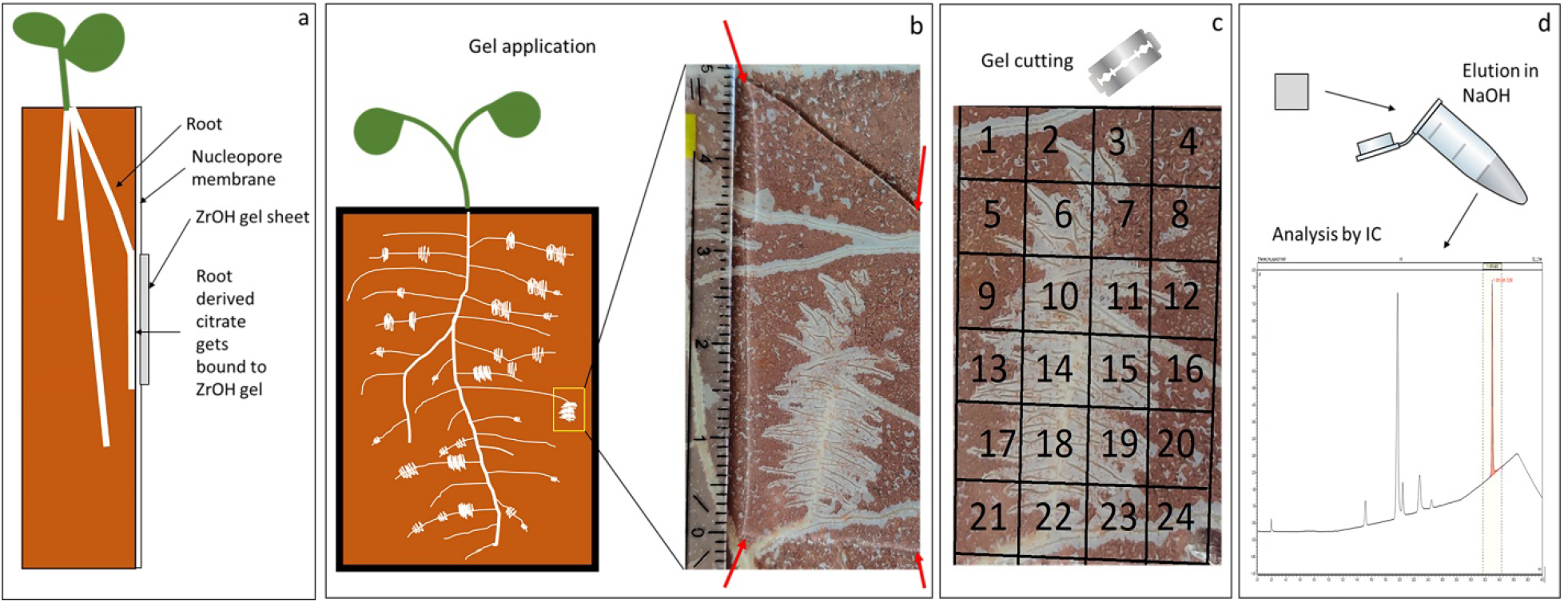
Schematic representation of citrate sampling with ZrOH gels of rhizotron grown plants. (a) Rhizotron grown plant. (b) ZrOH gel application on root/rhizosphere. (c) Gel cutting. (d) Citrate elution and IC analysis of eluates.

A small amount of water was added to the target root area until a thin film of water was visible at the polycarbonate membranes surface. The ZrOH gels were cut to the desired size depending on the selected sampling area and applied carefully on top of the polycarbonate membrane. A plastic foil was laid on top of the ZrOH gel to protect it against the back plate and the rhizotron was closed again. The gels were deployed for 24 hours. After the deployment they were placed on a clean and sterile (sterilisation with 3 mmol L^−1^ NaN_3_) piece of plastic foil and put in a clean and sterile airtight plastic bag together with a few drops of 3 mmol L^−1^ NaN_3_ to prevent microbial growth and possible citrate degradation. The plastic bags were stored at 4°C until cutting and citrate analysis. For citrate mapping the gels were cut with a clean and sterile razorblade into 5×5 mm pieces, 5×2 mm pieces and strips of 4- or 5-mm width, respectively. Every gel piece was eluted individually as described above and analysed for citrate.

### Citrate analysis

Citrate concentration was analysed by ion chromatography (IC) with suppressed conductivity detection (ICS-5000+ ion chromatographic system; AS-AP autosampler; ATC-3 trap column installed between pump and injection valve; IonPac 4×250 mm AS11-HC ion-exchange analytical column coupled with a AG11-HC 4mm guard column; anion self-regenerating suppressor ASRS 300 (4mm) in auto suppression recycle mode; Dionex, Thermo Fischer Scientific, USA). Ten µL sample was injected. Separation was performed with NaOH in laboratory grade I water as eluent, with a flow rate of 1.5 mL min^−1^ and constant temperature of 30°C, with the following gradient: E1 (NaOH 1mM) and E2 (NaOH 60mM); 0-15 min 20% E1 80% H_2_O, 15-25 min 15% E2 85% H_2_O, 25-35 min 30% E2 70% H_2_O, 35-45 min 60% E2 40% H_2_O. An external standard of citrate (Sigma Aldrich, citric acid, ACS reagent, ≥ 99.5%) was used for identification (retention time) and quantification after the construction of a calibration curve (R^2^=0.99).

## Results

### Characterization of ZrOH gel for citrate sampling

#### Citrate elution efficiency and uptake kinetics

Citrate elution efficiency parameters are shown in Table 1. The most efficient elution of citrate from the ZrOH gel disc (volume: 0.196 cm^3^) was accomplished using 1 mL 0.5 mol L^−1^ NaOH for 24 hours (Table 1) per gel disc. With these parameters the elution efficiency was found to be 89% ± 1.9%. Fig. 2A shows the citrate uptake kinetics on the ZrOH gel disc. In the first 2 hours the uptake was linear, binding up to 22.7 µg (0.11 µmol; 81.5%) of the citrate in the immersion solution. The remaining citrate was bound to the ZrOH gel after five hours.

**Figure 2.**
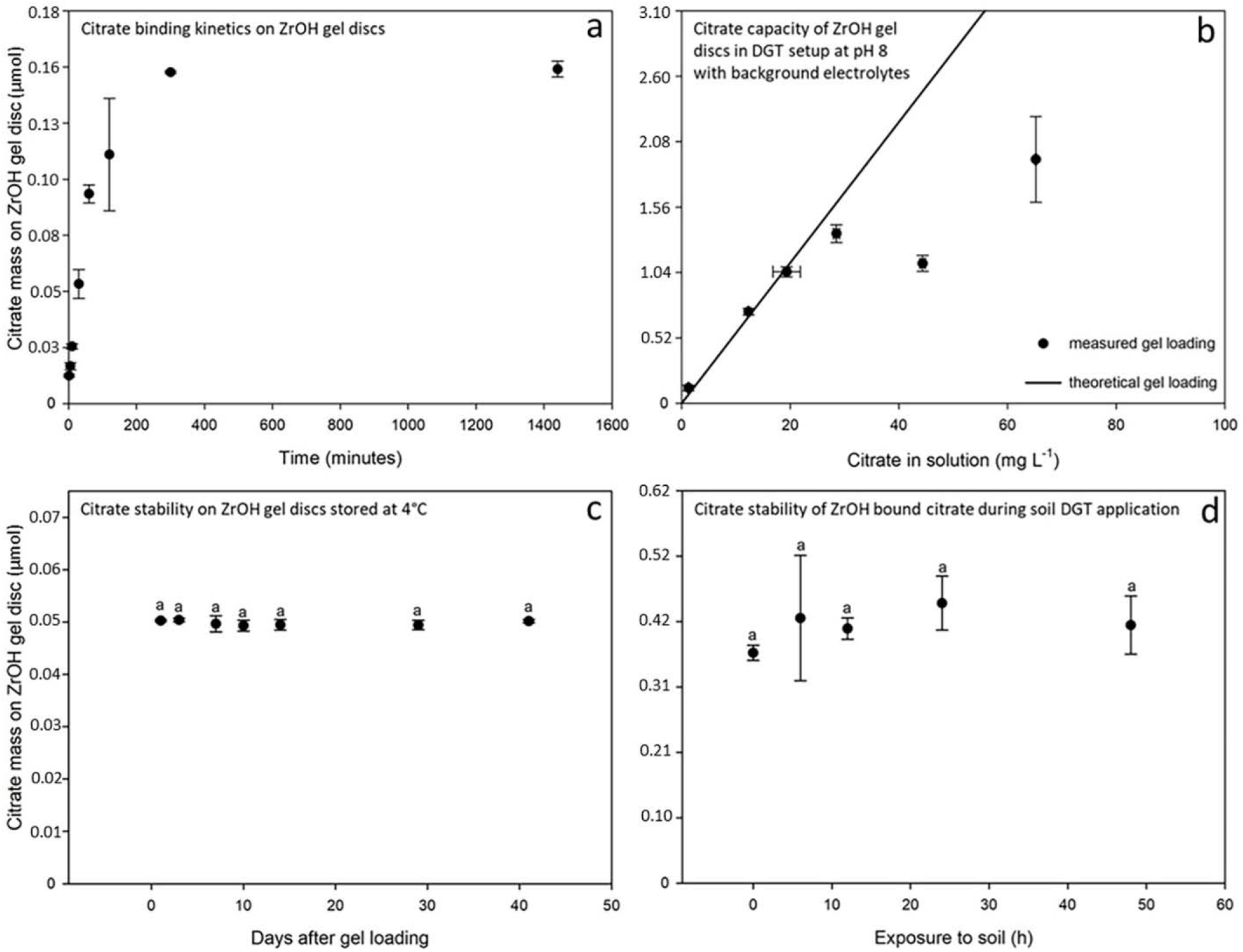
Chemical characteristics of ZrOH gels for citrate elution. (a) Citrate binding kinetics on ZrOH gels. (b) Representative citrate capacity of ZrOH gel discs in DGT setup at pH 8 with background electrolytes. (c) Citrate stability on ZrOH gel discs at 4°C. (d) Citrate stability of ZrOH bound citrate during soil DGT application. Letters indicate statistically significant difference according to one-way ANOVA comparing averages with the Holm Sidak post hoc test at p<0.05. Data points are means ± standard deviation of three replicates.

#### DGT binding capacity of precipitated ZrOH gel for citrate binding

Table 1 shows the DGT binding capacity of ZrOH discs for citrate uptake under infinite sink conditions. The experiment was performed at pH 4 and pH 8, either in laboratory water grade I or in a solution containing major inorganic soil solution anions and cations at a high, but realistic concentrations (ionic strength I = 15.8 mmol L^−1^). Fig. 2B shows a representative capacity graph for citrate. The capacity measured in solutions without background electrolytes was generally lower than the immersion solution containing the background electrolytes. Without background electrolytes, the DGT binding capacity was 0.63 µmol citrate gel disc^−1^ at pH 4, while the gel had a binding capacity of 0.16 µmol disc^−1^ at pH 8. In solutions containing background electrolytes, the DGT binding capacity was measured as 1.56 µmol gel disc^−1^ at pH 4 and as 1.04 µmol disc^−1^ at pH 8 (Fig S1).

#### Citrate stability on ZrOH gels and soil as potential citrate source

Exposure of citrate loaded ZrOH gels in the DGT setup to soil for up to 48 hours showed no significant alteration of the citrate mass bound to the ZrOH gel (Fig. 2D). No citrate was detected on unloaded ZrOH gels after exposure to unsterilized and sterilized bulk soil samples. Fig. 2C shows that no statistically significant citrate degradation occurred when citrate loaded ZrOH gels were stored for 41 days at 4 °C without NaN_3_.

### Case study: Mapping citrate exudation by *Lupinus albus* L. roots

Fig. 3 and Fig. S3 show citrate maps generated by deploying ZrOH gels on cluster roots of 45 days old white lupin plants. The citrate map depicted in Fig. 3A shows the exudation pattern of a white lupin cluster root of a plant which grew in the soil without P fertilization (0P). The map consists of 24 individual measurements performed on 5 × 5 mm gel pieces, into which the ZrOH gel was cut after citrate sampling. The cluster root was approximately 10 mm wide and 25 mm long and is representative of the cluster roots observed in the lupin root systems grown in this study. The majority of the citrate mass was observed at the very location of the cluster root, non-cluster root axes exuded almost no citrate (t-test, p = 0.002). After 24h of gel deployment, on average 0.06 µmol citrate cm^−2^ was detected in the gel pieces overlying parts of the cluster root. The highest measured gel loading was 0.16 µmol cm^−2^ for a cluster root area, whereas the lowest was 0.0004 µmol citrate cm^−2^, which was detected in a gel piece that was in contact with a soil containing no cluster root.

**Figure 3.**
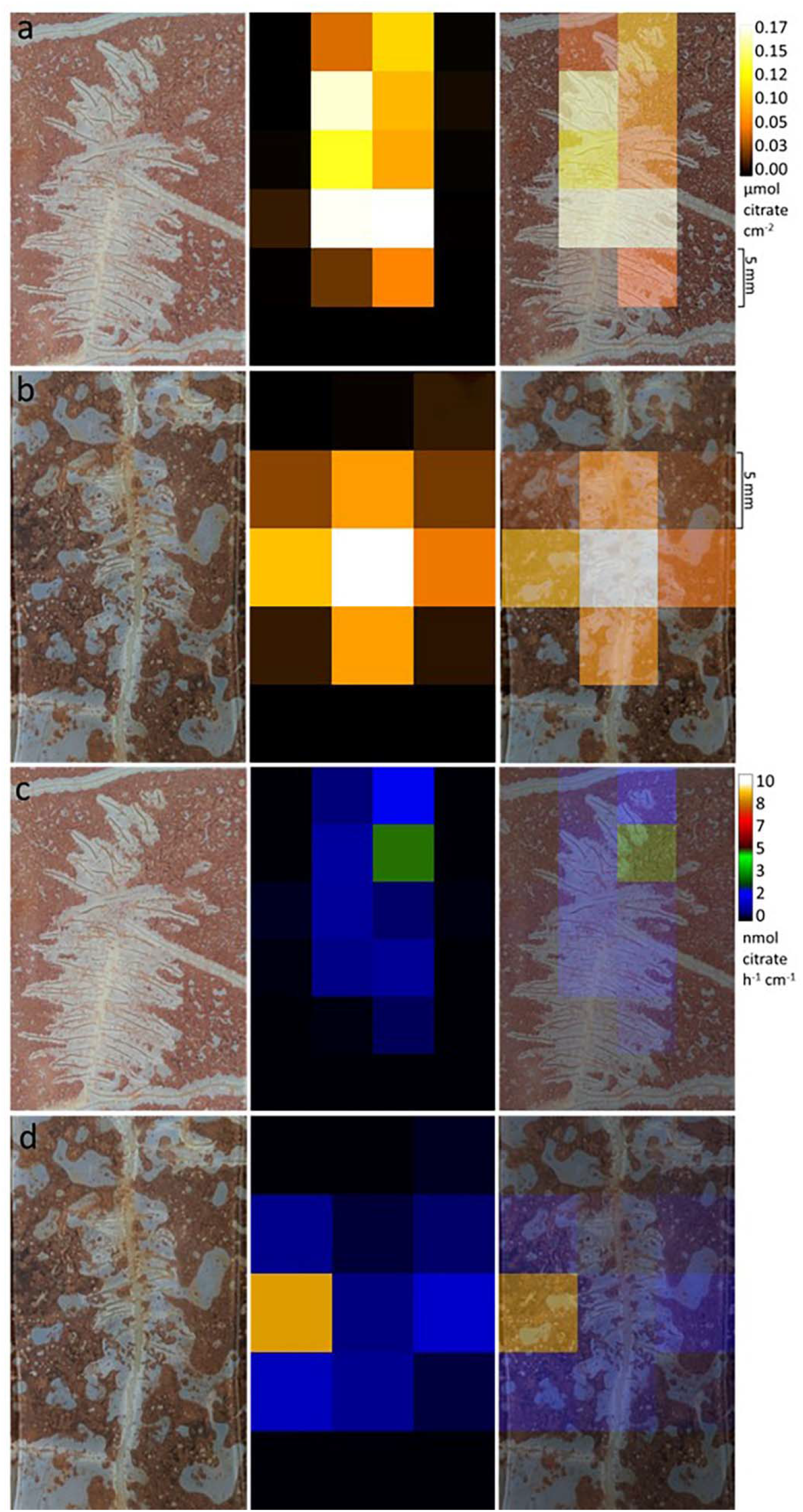
2D imaging of citrate exuded from two different white lupin cluster roots grown in unfertilized soil using the novel ZrOH sampling method. Squares have been cut into 5×5 mm pieces. a and b, exudation represented as µmol citrate cm^−2^ gel; c and d, exudation represented as µmol citrate h^−1^ cm^−1^ root length.

Fig. 3B shows a citrate map generated with 18 ZrOH gel pieces cut with a resolution of 5 × 5 mm. The representative cluster root used for the sampling was approximately 9 mm long and 5 mm wide and grew in a soil with 0P. The citrate map was very similar to the one shown in Fig. 3A. On average, 0.07 µmol citrate cm^−2^ were detected in the eluates of the gels in the cluster root region, whereas in non-cluster root gel pieces only 0.001 µmol citrate cm^−2^ were measured (t-test, p = 0.006). The smallest concentration measured in a gel piece was 0.003 µmol cm^−2^ at a non-cluster-root location, whereas the maximum amount was 0.16 µmol cm^−2^, exactly in the central part of the cluster root.

Fig. 4 shows citrate distributions of horizontally cut gels with 4 mm width. Fig. 4A shows a white lupin root grown in 250P soil. The gel was cut into 10 strips. A peak in citrate exudation was detected in the gel pieces overlying the cluster root, showing that the cluster root exuded significantly more citrate compared to non-cluster roots (0.31 nmol citrate in non-cluster roots *vs* 0.83 nmol citrate in cluster roots; t-test, p<0.001).

**Figure 4.**
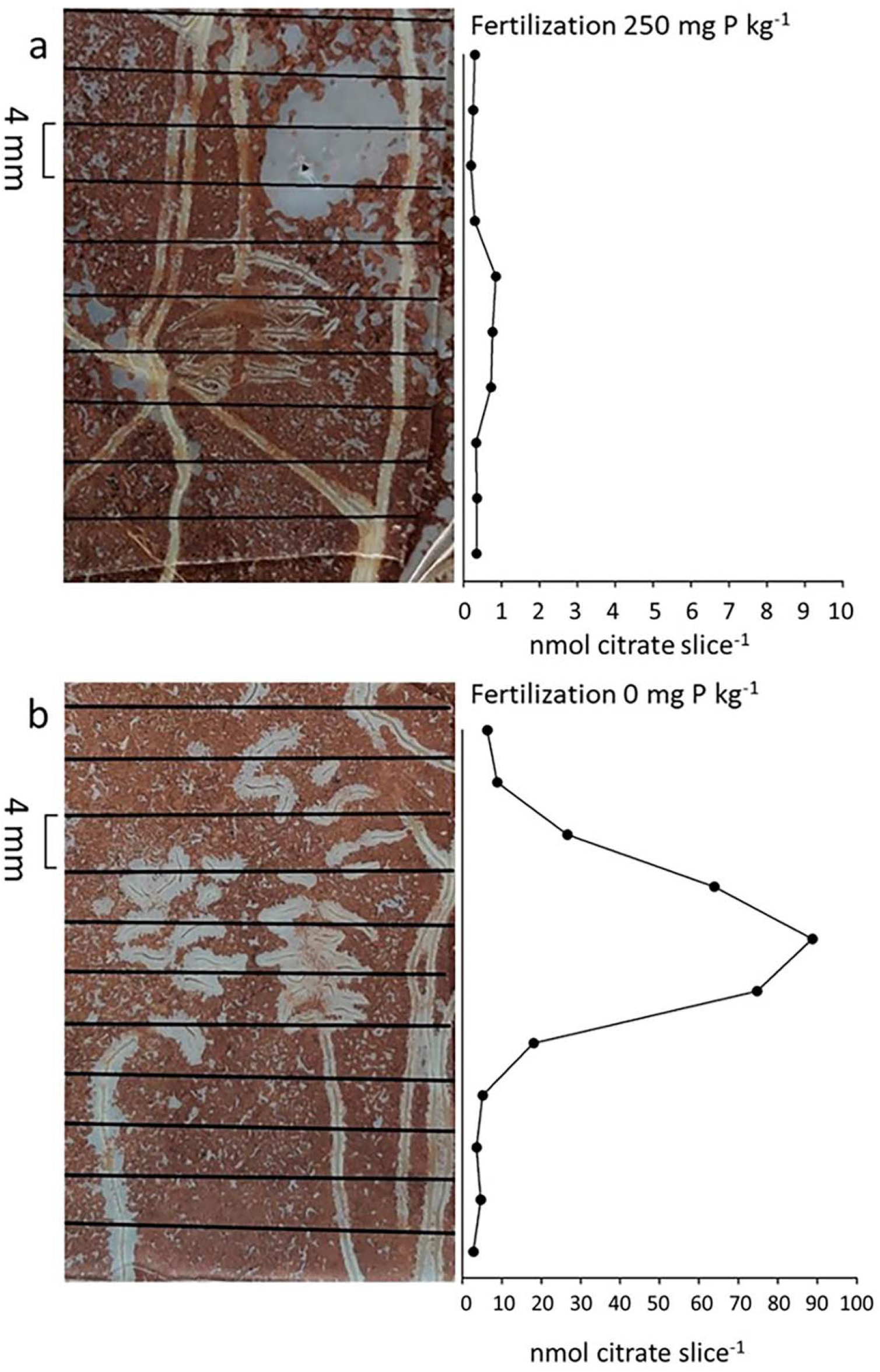
1D Profiles of citrate released from white lupin cluster roots. Gels were cut into 4 mm wide slices. Plants grown in a soil fertilized with a) 250 mg P kg^−1^ and b) 0 mg P kg^−1^. Note the different x-axis scaling: (a) 0-10 nmol cm slice^−1^, (b) 0-100 nmol slice^−1^.

The gel shown in Fig. 4B was deployed onto two cluster roots which grew in a 0P soil. The gel was cut into 11 strips. In the gel strips over the cluster roots were detected significantly higher amounts of citrate when compared with strips with non-cluster roots, 0.06 µmol citrate *vs* 0.001 µmol, respectively (t-test, p = 0.006). The pronounced citrate peak which can be seen in Fig. 4B corresponds to the cluster root.

## Discussion

### A straightforward and reliable DGT method for citrate sampling

In this study, ZrOH binding gel has been chosen as candidate for citrate sampling and was tested extensively for this purpose because of its known high capacity for the sorption of inorganic anions (Guan *et al*., 2015). The sorption kinetics shown in Fig. 2 confirms that citrate is adsorbed quantitatively and rapidly onto ZrOH gels. The stability/degradation tests showed that citrate was not released from gel discs once bound during 48-h sampling periods when citrate-loaded gels were in contact with unsterilized as well as with sterilized soil. Therefore, neither mineralization of citrate by microbes, nor competition for citrate by soil sorption sites reduced the amount of pre-loaded citrate on the DGT gel (Vives-Peris *et al*., 2019). These observations demonstrate that citrate bound to ZrOH gel was well protected during the sampling period. Moreover, citrate bound to the gel was not mineralized during gel storage at 4°C for up to 41 days, even without the addition of a biocidal chemical (*i*.*e*. NaN_3_, Fig. 2). This characteristic of citrate sampling by DGT represents a considerable advantage compared to other exudate sampling methods, as our method does not necessitate soil sterilisation to prevent mineralisation. Indeed, sterilisation procedures often introduce artefacts (Oburger and Jones, 2018).

The DGT capacity of the ZrOH gel for citrate depends on both the ionic strength and the pH of the soil solution (Table 1, Fig. S2). The acid dissociation constants of citrate are pK_a1_=3.13, pK_a2_=4.76 and pK_a3_=6.40 (Kraemer *et al*., 2006; Kettemann *et al*., 2016), while the point of zero charge of ZrOH has been reported to be pH_pzc_ 4.7 after hydration in distilled water for periods longer than about 4 days (Stanković *et al*., 2013). Our data showed a higher ZrOH capacity for citrate at pH 4 than 8. This decrease in the citrate capacity of ZrOH was expected given that the monovalent anion is the major citrate species at pH 4 and the net charge on ZrOH is slightly positive. On the contrary, at pH 8 the trivalent citrate anion is predominant and the net charge on ZrOH is negative. The DGT capacity of ZrOH for citrate was 0.63 µmol disc^−1^ (0.20 µmol cm^−2^) at pH 4 and 0.16 µmol disc^−1^ (0.05 µmol cm^−2^) at pH 8. The addition of background electrolytes at high, but realistic soil solution concentrations resulted in strongly increased DGT capacities: 1.56 µmol disc^−1^ (0.49 µmol cm^−2^) at pH 4 and 1.04 µmol disc^−1^ (0.33 µmol cm^−2^) at pH 8, respectively (Table 1). As a reference the capacity to bind PO_4_^3−^ anions in these gels is 1.72 µmol disc^−1^ (0.35 µmol cm^−2^) (Guan *et al*., 2015). We assume that mainly the multivalent cations (Ca^2+^, Mg^2+^, and the small concentration of Al^3+^) were responsible for this increase in citrate binding, as (1) cation sorption on ZrOH increases positive charges locally and also renders the net charge more positive, and as (2) these cations might act as cation bridges in binding citrate to ZrOH. Generally, the observed binding behaviour suggest electrostatic binding of citrate to the ZrOH phase in the gels used here.

Given these citrate binding properties, it is important to determine the soil pH and the soil solution composition before sampling citrate with ZrOH gels. The capacity values observed here are thresholds above which the gel ceases to take up citrate immediately and completely, *i*.*e*. where citrate binding ceases to conform to Eqn. (1). If citrate quantities larger than the DGT capacity are observed, the gel application period can be reduced to remain below capacity thresholds. However, in many cases, the concentration of root exudates is rather low (Vives-Peris *et al*., 2019; Wang and Lambers, 2019), thus exceedance of the capacity of the ZrOH gels should be very rare.

After the citrate adsorption on the gel it needs to be eluted for analysis. The elution procedure is easy to perform with high and well-reproducible elution efficiency (89 ± 1.9 %). We found that it is important to maintain a constant ratio of eluent volume to gel disc volume, *i*.*e*. 5 mL NaOH per cm^3^ of ZrOH gel (1 mL NaOH per ZrOH gel disc) in order to achieve this high elution efficiency reliably. The eluates can be stored at −20 °C. DGT pre-concentrates analytes on the gel and the resulting analyte concentrations in the eluate are often sufficiently high to allow analysis with conventional techniques, *e*.*g*. ion/liquid chromatography.

### Case study: Mapping citrate exudation by *Lupinus albus* L. roots

Fig. 3 shows, for the first time, 2D maps of citrate released by white lupin cluster roots grown in non-sterilized soil. This plant species is known to release huge amounts of citrate, yet to a different extent depending on the developmental stage of the cluster roots (*i*.*e*. juvenile, mature, senescent) (Shane and Lambers, 2005). The maximum citrate gel loading observed in this experiment was 0.16 µmol cm^−2^, which is well below the maximum DGT binding capacity at pH 5.2 and ionic strength I =15.8 mmol L^−1^ (as a measure of electrolytes) of the experimental soil used in this study, which we estimate at 0.31-0.47 µmol cm^−2^ given DGT binding capacity range determined at pH 4 and 8.

The exudation levels as well as the visual appearance of the two roots in Fig. 3 suggest that they are mature cluster roots (Uhde-Stone, 2017). Citrate exudation is a response to low soil P status in order to mobilize P. This is clearly shown in Fig. 4, as citrate exudation in P-fertilized soil is about 100-fold lower than in the unfertilized soil. The exudation maps show that the highest mass of citrate is released in the centre of the cluster root, where the root length density is also highest, translating to root-length based exudation rates being more homogeneously distributed across the cluster.

Citrate exudation represented as gel loading is hard to relate to citrate exudation rates based on root length or root mass, therefore the derivation of an exudation rate estimate from DGT data is desirable. While the DGT sampling setup and geometry do not allow for a direct calculation of exudation rates, an ‘apparent exudation rate’ can be determined by normalizing the amount of citrate measured per gel piece by the root length that this gel piece was in contact with and dividing by the deployment time (24 h). Given that the binding gel and the root were only separated by a 10-µm thin membrane layer, and that citrate is bound quickly and protected effectively from mineralization (Fig. **2D**), we assume that the predominant fraction of the citrate exuded by roots underlying the DGT gel is quantified. Apparent citrate exudation rates are shown in Fig. **3C,D**. Average apparent exudation rates (± standard deviation), calculated using only gel pieces in contact with cluster roots, are 0.92 ± 1.32 nmol cm^−1^ h^−1^ (n=13; Fig. **3C**) and 1.91 ± 3.22 nmol cm^−1^ h^−1^ (n=6; Fig. **3D**). These very high standard deviations are due to one individual very high apparent exudation rate value in each image, caused by a considerable amount of citrate detected at a location where root length was zero or very low. If these very high values are omitted, apparent exudation rate averages are 0.58 ± 0.57 nmol cm^−1^ h^−1^ (n=12; Fig. **3C**) and 0.61 ± 0.49 nmol cm^−1^ h^−1^ (n=5; Fig. **3D**).

The fraction of the citrate exuded directly into the soil from the root surface averted from the gel that is included in the obtained measurement is unclear, as a large part of this citrate amount is likely mineralized in the soil and does not diffuse towards the binding gel. However, assuming that the cluster roots exuded homogeneously around their circumference, the apparent exudation rate likely represents ∼50% of the total citrate exudation of the investigated rootlets. Based on the additional assumption that half of the cluster rootlets developed at the surface of the soil and thus contributed to DGT sampling, an estimate of the exudation rate of an entire cluster can be derived by multiplying the apparent exudation rate of the entire cluster by factor 4. Doing so, whole-cluster exudation rates of 44.3 nmol h^−1^ (Fig. **3C**) and 45.8 nmol h^−1^ (Fig. **3D**) can be derived. A comparison with published exudation rates of lupin cluster roots shows that these estimates agree very well with the range of measured values in hydroponic studies (0.25-11 nmol cluster^−1^ h^−1^, Table 2), showing that DGT-based citrate sampling is a promising tool to estimate realistic exudation rates.

**Table 2.** Apparent and estimated citrate exudation rates of whole mature root clusters grown under P deficiency. The exudation rates are based on the fluxes of citrate to the DGT as explained in the Discussion. * Values has been re-calculated using a mass of mature cluster roots of 50 mg fresh weight (Tiziani *et al.*, 2020).

|  | Apparent whole cluster<br>citrate exudation rate<br>(nmol h<sup>-1</sup>) | Estimated/measured<br>whole cluster citrate<br>exudation rate<br>(nmol h<sup>-1</sup>) | Reference |
| --- | --- | --- | --- |
| Root 1 (Fig. 3a, 3b) | 11.1 | 44.3 | this study |
| Root 2 (Fig. 3c, 3d) | 11.5 | 45.8 | this study |
|  |  | ~53.2 * | (Wasaki *et al.*, 2005) |
|  |  | ~13.3 * | (Wang *et al.*, 2007) |
|  |  | ~117 * | (Tomasi *et al.*, 2009) |
|  |  | ~42.3 * | (Shen *et al.*, 2005) |
|  |  | ~12.5 * | (Hagström *et al.*, 2001) |
|  |  | ~0.25 | (Tiziani *et al.*, 2020) |

### Appraisal of DGT-based citrate sampling and mapping

As discussed above, only a fraction of the exuded citrate is bound by the DGT gel in this method. One aim in further developing DGT-based citrate sampling will be to investigate and increase the accuracy of estimating cluster root exudation rates. In addition to more detailed experimental work, numerical simulation of root exudation and DGT sampling similar to the model used by Santner et al., (2012) who used a numerical model to study how phosphate released by roots is distributed on DGT gels, is a promising tool to increase the accuracy of estimating exudation rates. Adopting their approach should be uncomplicated for straight, individual root sections, but more complicated for the complex geometry of cluster roots.

Several techniques have been developed for collecting exudate at distinct root locations, including exudate sampling using filter papers (Neumann et al., 2014), agar gels (Hoffland et al., 1989), anion exchange membranes (Shi *et al*., 2011*a*) or micro suction cups (Dessureault-Rompré et al., 2006). Many of these techniques are, however, susceptible for sampling artefacts. For example, micro suction cups might alter local soil-chemical equilibria due to the continuous extraction of the soil solution. Agar based techniques have to be performed preferably with hydroponically grown plants and many other techniques disturb the roots by physical stress or artificial growing media (Vranová *et al*., 2013; Wang and Lambers, 2019). In comparison, DGT-based sampling is low-invasive, protects the exudate sample from mineralization once it is bound to the binding material in the gel, and operates in non-sterilized soil. Moreover, it is the only technique offering the possibility to generate exudate maps at millimetre resolution.

DGT-based citrate sampling can be utilized for studying the exudation behaviour of plant roots in the soil environment at unprecedented spatial resolution. Successive replacement of the DGT binding gel should allow for obtaining information on the time-course of exudation. Provided that suitable binding phases are available, the DGT sampling principle can be easily adopted for sampling a large range of exudate compounds such as other organic acids, amino acids, phenols and carbohydrates. The focus of this study was on resolving the spatial distribution of exudation. However, it is also possible to elute large gel sheets that cover representative parts of the root system of crop plants. This approach could allow for screening at least small sets of cultivars or species for desired root exudation traits like high exudation rates or the occurrence of specific exudate compounds, and in this way might be useful in cultivar selection and breeding.

## Supplementary data

Materials and methods: Description of the soil liming and fertilization procedure, as well as the description of the determination of the citrate diffusion coefficient in diffusive gel

Results of the diffusion coefficients and a 2D imaging of a cluster root

Fig. S1. Capacity plots of citrate for ZrOH gels

Fig. S2. Citrate diffusion experiments plots

Fig. S3. Citrate mapping of a cluster root

Table S1. Chemical characteristics of the soil used in the plant experiments.

## Acknowledgement

This work was supported by grants from the Austrian Science Fund (FWF) and the Federal State of Lower Austria: P27571-BBL and Free University of Bolzano (NUMICS TN200E). Erik Smolders contributed to this work during a sabbatical leave to the University of Natural Resources and Life Sciences, Vienna (BOKU), supported by grant K801219N from FWO-Vlaanderen. We also thank Gerlinde Wieshammer, Sarah Mühlbacher, Federica De Berardinis, Stefan Wagner and Christina Roschitz for their help during the experiments, analysis and DGT preparation.

## Author contribution

RT, JS, MP, ES, TM designed the study; RT, JCH and ES performed the experiments; RT, TM, CS and JS wrote the manuscript with editorial input of all co-authors. All authors read and approved the manuscript.

## Data availability statement

The data supporting the findings of this study are available from the corresponding author, (Raphael Tiziani, Jakob Santner), upon request.

